# Dual TCR-alpha expression on MAIT cells as a potential confounder of TCR interpretation

**DOI:** 10.1101/2021.03.25.436871

**Authors:** Sara Suliman, Lars Kjer-Nielsen, Sarah K. Iwany, Kattya Lopez Tamara, Liyen Loh, Ludivine Grzelak, Katherine Kedzierska, Tonatiuh A. Ocampo, Alexandra J. Corbett, James McCluskey, Jamie Rossjohn, Segundo R León, Roger Calderon, Leonid Lecca Garcia, Megan B. Murray, D. Branch Moody, Ildiko Van Rhijn

## Abstract

Mucosal-associated invariant T (MAIT) cells are innate-like T cells that are highly abundant in human blood and tissues. Most MAIT cells have an invariant T cell receptor (TCR) α chain that uses TRAV1-2 joined to TRAJ33/20/12 and recognize metabolites from bacterial riboflavin synthesis bound to the antigen-presenting molecule, MR1. Recently, our attempts to identify alternative MR1-presented antigens led to the discovery of rare MR1-restricted T cells with non-TRAV1-2 TCRs. Because altered antigen specificity is likely to lead to altered affinity for the most potent known antigen, 5-(2-oxopropylideneamino)-6-D-ribitylaminouracil (5-OP-RU), we performed bulk TCRα and β chain sequencing, and single cell-based paired TCR sequencing, on T cells that bound the MR1-5-OP-RU tetramer, but with differing intensities. Bulk sequencing showed that use of V genes other than TRAV1-2 was enriched among MR1-5-OP-RU tetramer^low^ cells. Whereas we initially interpreted these as diverse MR1-restricted TCRs, single cell TCR sequencing revealed that cells expressing atypical TCRα chains also co-expressed an invariant MAIT TCRα chain. Transfection of each non-TRAV1-2 TCRα chain with the TCRβ chain from the same cell demonstrated that the non-TRAV1-2 TCR did not bind the MR1-5-OP-RU tetramer. Thus, dual TCRα chain expression in human T cells and competition for the endogenous β chain explains the existence of some MR1-5-OP-RU tetramer^low^ T cells. The discovery of simultaneous expression of canonical and non-canonical TCRs on the same T cell means that claims of roles for non-TRAV1-2 TCR in MR1 response must be validated by TCR transfer-based confirmation of antigen specificity.

## Introduction

Adaptive cellular immunity relies on recombination of the T cell receptor (TCR)-β (TRB), TCR-γ (TRG), TCR-α and TCR-δ (TRA/TRD) genomic loci during T cell development in the thymus^1^. Remarkable TCR diversity is achieved by combinatorial usage of genome-encoded variable (V), diversity (D), and joining (J) genes, and addition of intervening non-templated (N) nucleotides^2^. Many T cells recognize peptide antigens in the context of highly polymorphic human leukocyte antigen (HLA) molecules^3^. In parallel, some T cells bind non-peptide antigens presented by non-MHC-encoded antigen-presenting molecules, including the MHC-related protein 1 (MR1) and cluster of differentiation (CD)1 proteins (reviewed in ^4, 5^). Unlike MHC, CD1 and MR1 proteins are almost monomorphic^6^, and consequently CD1- and MR1-reactive T cells tend to express characteristic TCR motifs, shared by many individuals irrespective of their HLA haplotypes^7^. These invariant TCR motifs^7^ recognize unique antigen classes, including pathogen-derived mycobacterial lipids for CD1b^8^, α-galactosyl ceramides for CD1d^9^ and metabolites from active bacterial biosynthetic enzymes for MR1^10^. These invariant TCRs are thought to have co-evolved with cognate nonclassical antigen-presenting molecules in different species^11^.

Due to their potential to elicit generalizable population-level immune responses, donor-unrestricted T cells (DURTs), and the antigens they recognize, are attractive targets of vaccination against microbes like *Mycobacterium tuberculosis* (*Mtb*)^12^. In particular, mucosal-associated invariant T (MAIT) cells, which recognize antigens presented by MR1, are attractive candidates due to their abundance in the blood^13^, their high reactivity against several bacterial infections^14, 15, 16, 17^, and their documented roles in vaccination^18, 19^. MR1 tetramers bind directly to TCRs and allow for unequivocal identification of MAIT cells and more diverse MR1-restricted αβ^20^ and γδ^21^ T cells, and provide a unique opportunity to identify novel TCR rearrangements and antigen specificities^22^. Human MAIT TCRα chains display a characteristic complementarity-determining region (CDR3α) formed by a rearrangement between TRAV1-2 and TRAJ33, or sometimes TRAJ12 or TRAJ20, with few non-template encoded (N)-nucleotides^22, 23, 24^, and a biased preference for some TRB genes^23, 25, 26^. Diversity in TRB gene usage in MAIT cells is potentially associated with recognition of different microbes^25, 27, 28, 29^ or different ligands^30^. These canonical MAIT cells have a preferred specificity for 5-(2-oxopropylideneamino)-6-D-ribitylaminouracil (5-OP-RU) over 6-formylpterin (6-FP)^10, 31, 32^. Whereas TCR conservation, especially ‘canonical’ TRAV1-2 usage has been considered a key defining feature of human MAIT cells for decades, a new direction in the field has resulted from identification of ‘non-canonical’ TRAV1-2-negative (TRAV1-2^−^) and γδ T cells^21^ that recognize MR1 and suggested to have unique antigen specificities^20, 33, 34, 35, 36, 37^. MAIT cells have broadly reported roles in infection^17^, cancer^38^, and autoimmunity^39^. Hence, defining MAIT TCR motifs can be used to infer pathogenic and protective TCR clonotypes relevant to immunodiagnosis or vaccination.

Several new technologies and algorithms for high-dimensional TCR sequencing analysis have successfully identified clonally expanded populations of antigen-specific T cells, and their TCR motifs among large numbers of blood- and tissue-derived T-cells^40, 41, 42, 43^. These sequencing technologies derive TCR sequences either from single cells, which identify paired TCRα and TCRβ^44, 45^, or bulk genomic^46^ or transcriptomic sequencing data^41, 47^. In this study, we sought to use MR1 tetramers and high throughput TCR sequencing to identify non-canonical TCR patterns. We observed MAIT cell populations with differing binding intensities to the 5-OP-RU-loaded MR1 tetramers. We hypothesized that MAIT cells with lower MR1-tetramer binding intensities would reveal unique TCR motifs consistent with lower preferential binding to the 5-OP-RU/MR1 antigen complex. Consistently, we detected an enrichment of TRAV1-2^−^ TCRs in MR1 tetramer^+^ MAIT cells, especially those with lower MR1-tetramer intensity. However, detailed TCR gene transfer studies revealed that the lower tetramer binding was explained by dual expression of canonical and non-canonical TCRα chains in the same TRAV1-2^+^ clonally expanded MAIT cells, as opposed to a single non-canonical TCR with lower affinity for MR1-5-OP-RU. Dual TCR expression previously observed in HLA-restricted^48^ and CD1d-reactive T cells^49^, but takes on special importance in the MAIT cell system because it can confound the assignment of non-canonical TCRs for MR1 specificity. These data establish the need to validate the antigen specificity of newly-described TCR motifs from large-dimensional sequencing platforms by TCR gene transfer and other alternative techniques^50^.

## Materials and Methods

### Human participants

#### Lima, Peru

We recruited Peruvian participants with active TB disease, or asymptomatic household contacts of TB cases with positive or negative QuantiFERON TB Gold-In-tube test results from Lima, Peru, as described previously^51, 52^. The Institutional Review Board of the Harvard Faculty of Medicine and Partners HealthCare (protocol number IRB16-1173), and the Institutional Committee of Ethics in Research of the Peruvian Institutes of Health approved this study protocol. All adult study participants and parents and/or legal guardians of minors provided informed consent, while minors provided assent. The protocol is approved by the Institutional Review Board of Harvard Faculty of Medicine and Partners HealthCare, and Institutional Committee of Ethics in Research of the Peruvian Institutes of Health.

#### Boston, USA

We obtained de-identified leukoreduction filters (leukopak) samples from healthy blood bank donors through the Brigham and Women’s Hospital Specimen Bank, as approved by the Institutional Review Board of Partners HealthCare.

#### Tennessee, Memphis

Peripheral blood mononuclear cell (PBMC) samples were obtained from healthy children, adult, and elderly donors from St. Jude Children’s Research Hospital (XPD12-089 IIBANK and 1545216.1).

#### Melbourne, Australia

Spleen (SP) lymphoid tissues were collected from deceased donors, whose mortality was caused by conditions other than influenza (DonateLife, Australia), after written informed consent was given by next of kin^53^. The University of Melbourne Human Ethics Committee approved experiments (identification numbers 1443389.4, 1955465 and 1545216.1).

### Flow cytometry analysis

The protocol and primary analysis of Peruvian samples by flow cytometry was reported previously^51^. MR1 monomers were obtained from The University of Melbourne, Australia^10, 22^, and used to generate tetramers in Boston as previously described^51^. For HEK293T cell validation experiments, we used MR1 tetramers obtained from the National Institutes of Health (NIH) Tetramer Core facility.

### Genomic bulk TCR sequencing (Adaptive Biotechnologies, Seattle)

For TCR sequencing from genomic templates, 3900 MR1 tetramer^hi^ and 4500 MR1 tetramer^int^ cells were doubly sorted from PBMC samples from Peruvian donor 58-1 after 14 days of polyclonal T cell expansion. For expansion, 10^6^ cells were cultured with 25 × 10^6^ irradiated allogeneic PBMC, 5 × 10^6^ irradiated allogeneic Epstein Barr Virus transformed B cells, 30 ng/ml anti-CD3 monoclonal antibody (clone OKT3) for 14-16 days, in the presence of 1 ng/ml interleukin-2 (IL-2)^52^. PBMC samples from healthy Boston blood bank donors LP1 and CO2 were not expanded before double cell sorting. Cell numbers obtained from the sorted tetramer^hi^, tetramer^int^, and tetramer^low^ populations were 2000, 5800, 3100, respectively, for LP1 and 1100, 4000, and 2300, respectively, for CO2. High-throughput TCR sequencing and assignment of V and J genes was performed for the TCRβ locus and the TCRαδ locus (Adaptive Biotechnologies, Seattle, WA) using a multiplex PCR approach on genomic DNA isolated from sorted T cells using the Qiagen QIAamp DNA Mini Kit, followed by Illumina high-throughput sequencing^46^.

### Sorted single cell paired TCR sequencing

Single-cell TCR sequencing was adapted from a previously published protocol^41^. Briefly, single MR1-tetramer-binding cells from Peruvian participant 7-3 and blood bank donors 702A and 703A were sorted into 96-well plate coated with Vapor-Lock (Qiagen) containing Iscript cDNA synthesis mixture (Bio-Rad) and 0.1% triton X-100 for direct cell lysis. Reverse transcription was performed in a thermocycler (25°C for 5’, 42°C for 30’, 80°C for 5’). Subsequently, cDNA samples were amplified in a nested PCR reaction using Denville Choice Taq Polymerase (Thomas Scientific), using previously described primers^41^. Briefly, the first external reaction contained a mixture of all TCRα and TCRβ forward primers, combined at 1 μM each, and reverse TRAC and TRBC primers at 10 μM each: 95°C for 2’, 35 cycles of (95°C for 20’’, 50°C for 20’’, 72°C for 45’’), and 72°C for 7’. A second internal PCR reaction used a mix of TCRα forward primers at 1 μM each with a reverse internal TRAC primer at 10 μM, or a mix of TCRβ forward primers and reverse TRBC primer, separately at cycling conditions: 95°C for 2’, 35 cycles of (95°C for 20’’, 56°C for 20’’, 72°C for 45’’), and 72°C for 7’ using previously described primers^41^. Amplicons were analyzed on an agarose gel, and bands were excised using a UV lamp and purified using the QIAquick Gel Extraction Kit (Qiagen) then sent for Sanger sequencing (Genewiz). Sequences were reverse-complemented and analyzed using 4Peaks software and mapped to the reference sequences for the genome-encoded V and J segments for both the TCRα and TCRβ genes on the ImMunoGeneTics (IMGT) information system database. The unmapped sequences were considered N-nucleotides, and/or Dβ segments for TCRβ to determine the complementarity-determining region (CDR)-3. CDR3α and CDR3β amino acid sequences were predicted by in silico translation, showing productive in-frame rearrangements, using the online ExPASy translate tool (https://web.expasy.org/translate/).

For Australian samples, single MR1-5-OP-RU-tetramer^+^TRAV1-2^+^ PBMCs from healthy donors and spleen tissues were sorted using a FACSAria (BD Biosciences) into 96-well plates. Paired CDR3αβ regions were determined using multiplex-nested reverse transcriptase PCR before sequencing of TCRα and TCRβ products, as previously described^41, 54^, and reported^55^. For paired TCRαβ analyses, sequences were parsed into the IMGT/HighV-QUEST web-based tool using TCRBlast1 (kindly provided by Paul Thomas and Matthew Caverley), to determine V(D)J regions.

### TCR transfection assay

Synthetic TCRα and TCRβ sequences (Genewiz) from MR1 tetramer-binding sorted single T cells, separated by self-cleaving Picornavirus 2A (P2A)-linker sequence (GGATCCGGCGCCACCAATTTCTCGCTGCTTAAGCAGGCCGGCGACGTCGAAGAGAACCCCGGGCCC**ATG**), were cloned into a GFP-containing pMIG vector using standard restriction digestion and cloning procedures. Human embryonic kidney (HEK293T) cells were cultured overnight on a 6-well plate containing 4 mL of Dulbecco’s Modified Eagle Medium-10 media supplemented with 10% fetal bovine serum and penicillin-streptomycin at 37°C, and were subsequently co-transfected with the pMIG-TCR and pMIG-CD3 plasmid^56^ using FuGENE HD transfection reagent (Promega). Transfected HEK293-T cells were analyzed for tetramer binding by flow cytometry 48-72 hours following transfection. Antibodies used to stain transfected 293T cells were Brilliant Violet 421-conjugated anti-human CD3 antibody (Biolegend) and phycoerythrin (PE)-conjugated anti-human TCRαβ antibody (BD BioSciences).

## Results

During a quantitative study of MAIT cells in a Peruvian TB cohort^51^, we observed MAIT cell populations with variable staining intensities for the 5-OP-RU-loaded MR1 tetramer **(Figure 1A)**. This phenomenon was observed in participants with and without evidence for *Mtb* infection and did not seem to be correlated with TB disease. Whereas canonical MAIT TCRs typically show high affinity for MR1-5-OP-RU, we hypothesized that MAIT cells with lower tetramer staining intensity may reflect different and variable TCR motifs, consistent with their lower affinities to the MR1-5-OP-RU complex. To define TCR gene usage in high, intermediate and low staining populations, we sorted MAIT cell populations with different MR1-tetramer staining intensities and performed bulk TCRα and TCRβ sequencing from genomic DNA and subsequent V- and J-gene assignment of rearranged genes. Subsequently, we sorted MAIT cell populations from one Peruvian sample (58-1) after polyclonal T cell expansion, and from two random Boston blood bank donors (LP1 and CO2) without expansion **(Figure 1B)**. The populations were sorted based on MR1-tetramer fluorescence intensities and re-sorted prior to sequencing to ensure purity and preservation of MR1-tetramer binding levels **(Supplementary Figure 1)**. Regardless of the source of PBMCs, we saw similar patterns with TRAV1-2 TCRs in brightly staining cells, and TCRα V-genes other than Vα7.2 (TRAV1-2) were enriched in sorted MAIT populations with low and intermediate MR1-tetramer staining **(Figure 1B-C).** This pattern of atypical TRAV gene usage in MAIT cells with lower MR1-tetramer binding relative to MAIT cells with high MR1-tetramer staining was observed even after discarding unproductive TCRα chains **(Supplementary Table 1)**. Frequencies of TRAV1-2^−^ MAIT cells in blood did not differ by TB status in Peruvian samples ( Kruskal-Wallis: *p=0.*75; **Figure 1D)**. TRAV1-2^−^ MAIT in these samples **(Figure 1B-C)** were similar to frequencies previously reported in other populations^20^ representing a minority of T cells (0.6-40%) but they were potentially biologically significant because TCRα diversity diverges from the conventional understanding of MAIT cell function.

**Figure 1:**
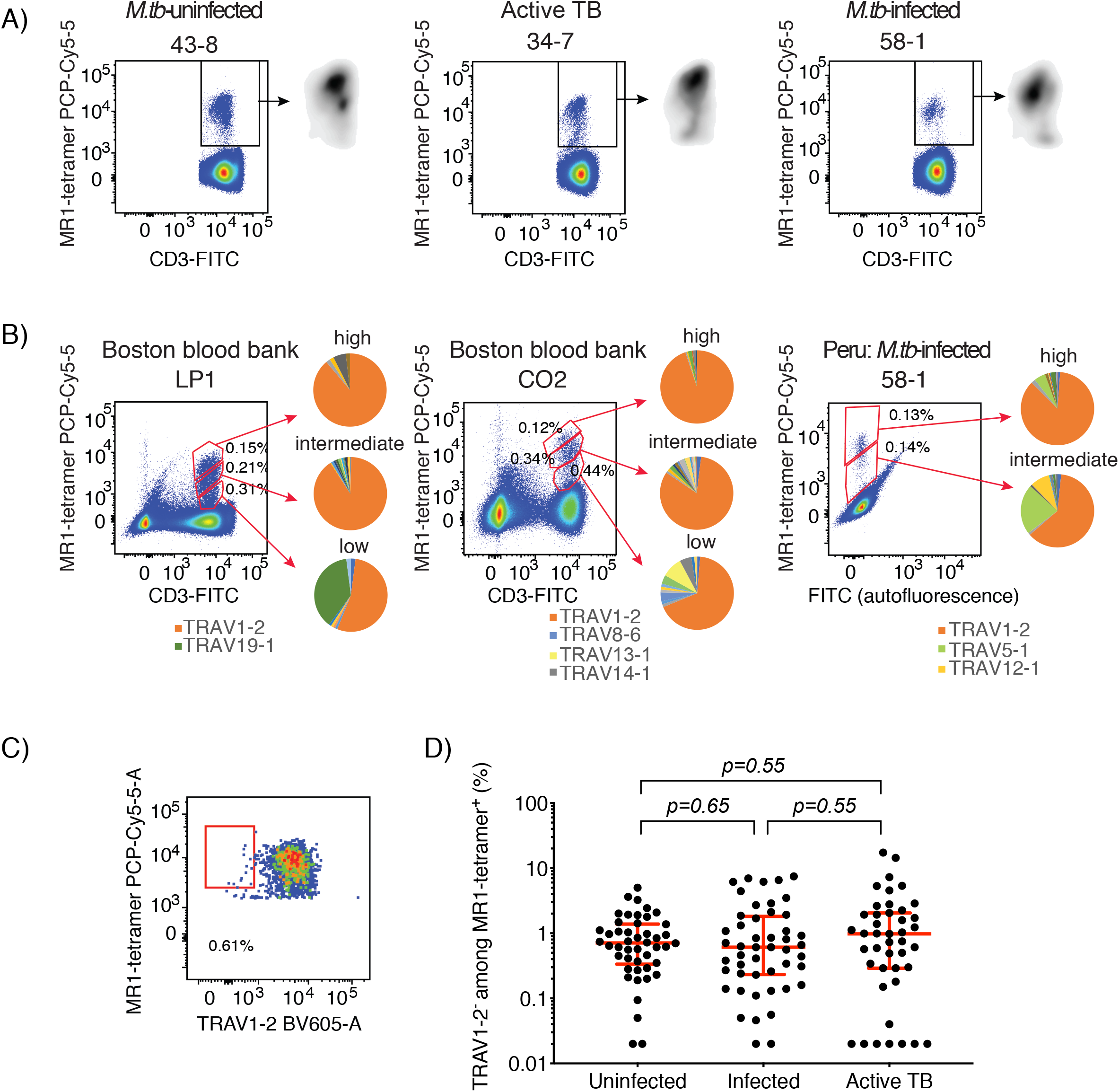
TRAV1-2^−^ TCR sequences are enriched in MAIT cells with lower MR1 tetramer staining intensities. (A) Three examples of variable MR1 tetramer staining intensities by flow cytometry in pre-gated T lymphocytes in samples from uninfected, latent, and active TB participants. (B) Gating strategy for bulk-sorted MAIT cells with different 5-OP-RU-loaded MR1 tetramer staining intensities is shown. The pie charts depict distribution of TCRα gene usage from the different populations. (C) Gating strategy to identify TRAV1-2^−^ MAIT cells among all MR1-tetramer-binding cells is shown. (D) Proportions of TRAV1-2^−^ MAIT cells among all MR1-tetramer-binding cells in the Peruvian samples from healthy participants who are either uninfected or infected with *Mycobacterium tuberculosis*, and active TB patients are shown. Error bars denote medians and interquartile ranges.

We sought to explain the discrepancy between the low frequencies of TRAV1-2^−^ MAIT cells as determined by flow cytometry **(Figure 1D)**, and the higher frequencies of TRAV1-2^−^ TCRα chain sequences identified in sorted MAIT cells, as determined by bulk TCR sequencing **(Figure 1C).** Hence, we sorted single cells from populations with different MR1-tetramer binding levels from one Peruvian participant, where we detected three clear MR1-tetramer binding levels (MR1-tetramer^high^, MR1-tetramer^int^, and MR1-tetramer^low^), and applied a previously described nested PCR protocol to cDNA amplified from each single cell^41^ to determine the sequences of paired TCRα and TCRβ chains **(Figure 2A)**. Non-TRAV1-2 TCRα gene usage was enriched in populations with lower MR1-tetramer binding, with 15/40 (37.5%) of the MR1-tetramer^int^ cells using TRAV16 and identical CDR3α nucleotide sequences, and 14/34 (41.2%) of the MR1-tetramer^low^ cells using TRAV5, of which 13 had identical CDR3α nucleotide sequences, suggesting clonal expansion *in vivo* **(Figure 2A and Supplementary Tables 2 and 3)**. Similarly, we detected TRAV1-2^−^ TCRs from single cell-sorted MR1-tetramer^low^ populations from two healthy blood bank donors: 1/33 (3%) and 8/48 (16.7%), but none in MR1-tetramer^high^ counterparts **(Figure 2B)**. Furthermore, the atypical TRAJ33^−^ joining regions were seen more frequently in low MR1 tetramer staining cells. Overall, these patterns from oligoclonal T cells (**Figure 2**) matched those of polyclonal T cells (**Figure 1**) and demonstrated more non-canonical gene usage in TCRs among low MR1 tetramer staining T cells.

**Figure 2:**
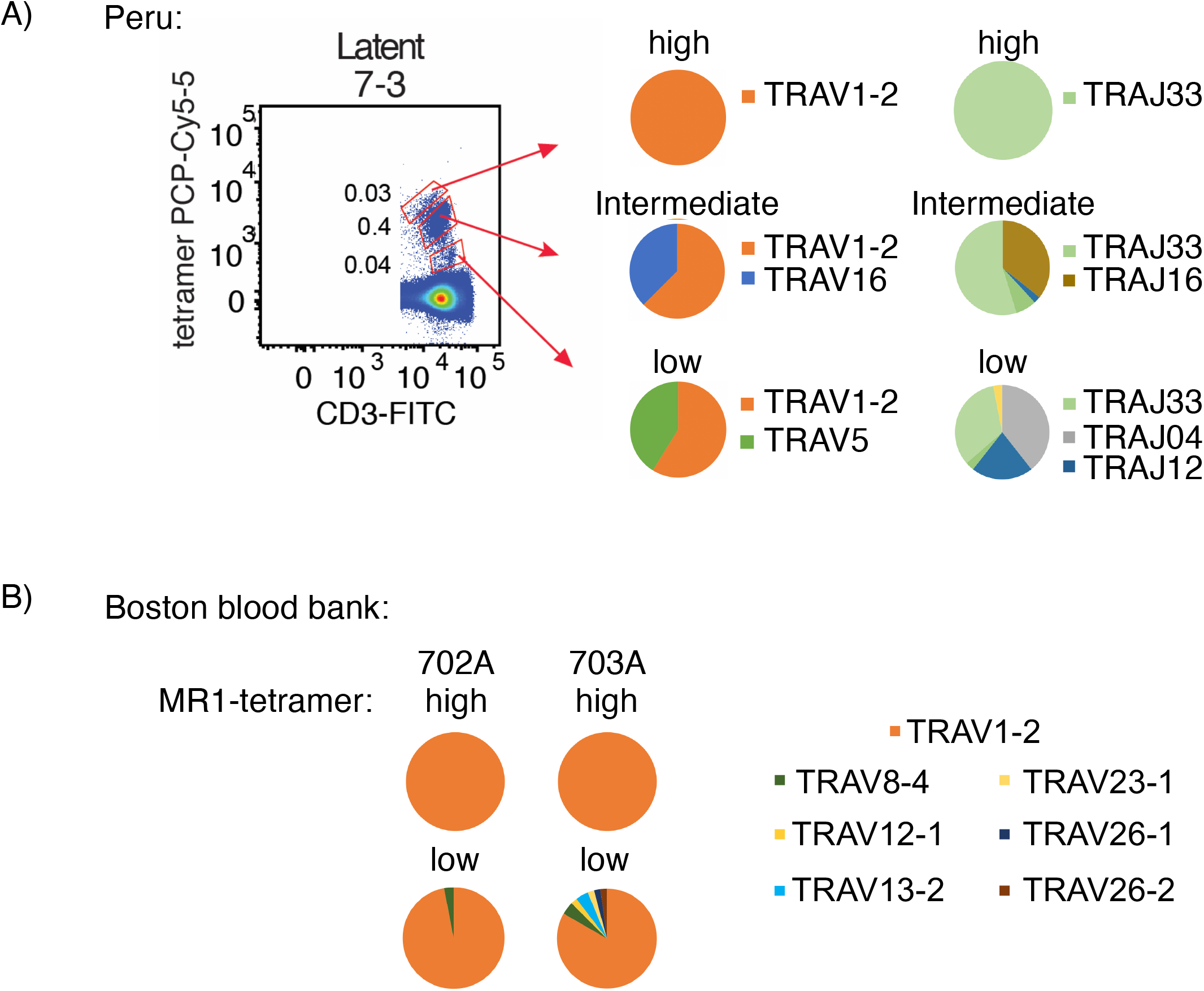
Single-cell sorted MAIT cells also show enrichment of TRAV1-2-negative TCR sequences. (A) Gating strategy shows single cell-sorted MAIT cells with different 5-OP-RU-loaded MR1 tetramer staining intensities in Peruvian latent sample no. 7-3. The pie charts depict distribution of TCRα gene usage from the different sorted populations. (B) Pie charts showing distribution of TCRα V-gene usage in single-cell-sorted MR1-tetramer^high^ and MR1-tetramer^low^ T cells from two additional healthy blood bank donors.

To validate the MR1-reactivity of these putative MAIT TCRs, we co-transfected human embryonic kidney (HEK293T) cells with pMIG vectors expressing CD3 and the paired TCRα and TCRβ sequences derived from three clones with non-TRAV1-2 TCR sequences **(Figure 3)**, which showed clear clonal expansion in samples analyzed with bulk **(Figure 1B)** or single cell **(Figure 2A)** TCR sequencing methods. Next, we measured TCR binding to the 5-OP-RU-loaded MR1 tetramer **(Figure 3)**. We also transfected TCRα and TCRβ from a canonical MAIT TCR (TRAV1-2-TRAJ33) identified in the bulk-sorted MR1-tetramer^high^ cells as a positive control **(Figure 3)**. Co-transfected HEK293T cells co-expressed CD3 and TCRαβ on the cell surface **(Figure 4, left)**. The 5-OP-RU-loaded MR1-tetramer, but not the MR1 tetramer loaded with the non-agonist 6-FP-loaded MR1 tetramer, stained CD3^+^ cells from HEK293T cells transfected with the TRAV1-2^+^ TCR, as expected. However, the MR1 tetramers, loaded with either 6-FP or 5-OP-RU, did not bind cells expressing the TRAV1-2^−^ TCRs identified in MR1-tetramer^low^ and MR1-tetramer^int^ populations **(Figure 4)**, despite the original detection of these TCR sequences in MR1-tetramer-binding cells **(Figures 1 and 2)**.

**Figure 3:**
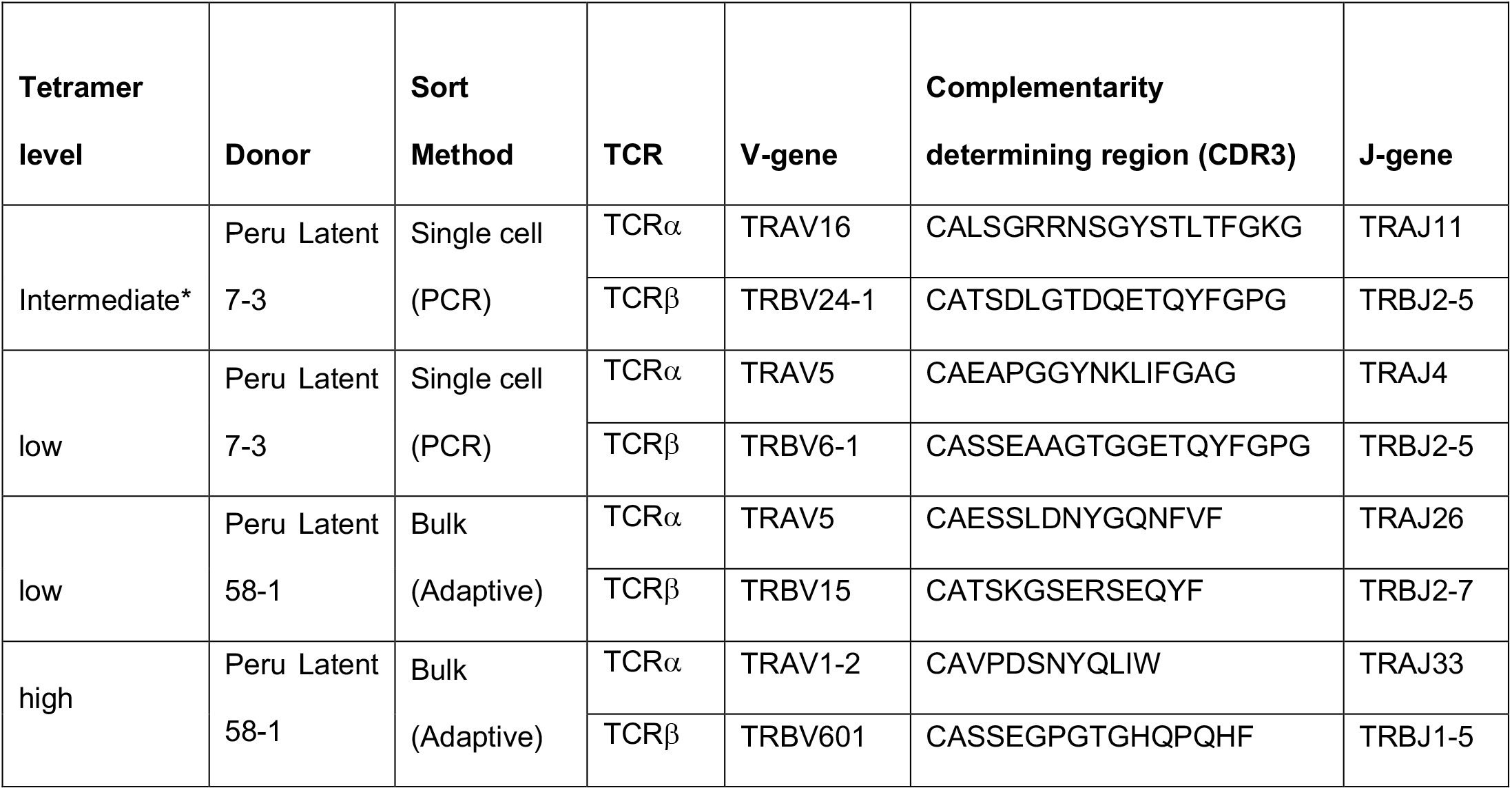
T-cell receptor sequences for additional validation by HEK293T cell transfection experiments. * Templates from this reaction were re-amplified using TRAV1 forward primer only with TRAC reverse primer **(Figure 4)**.

**Figure 4:**
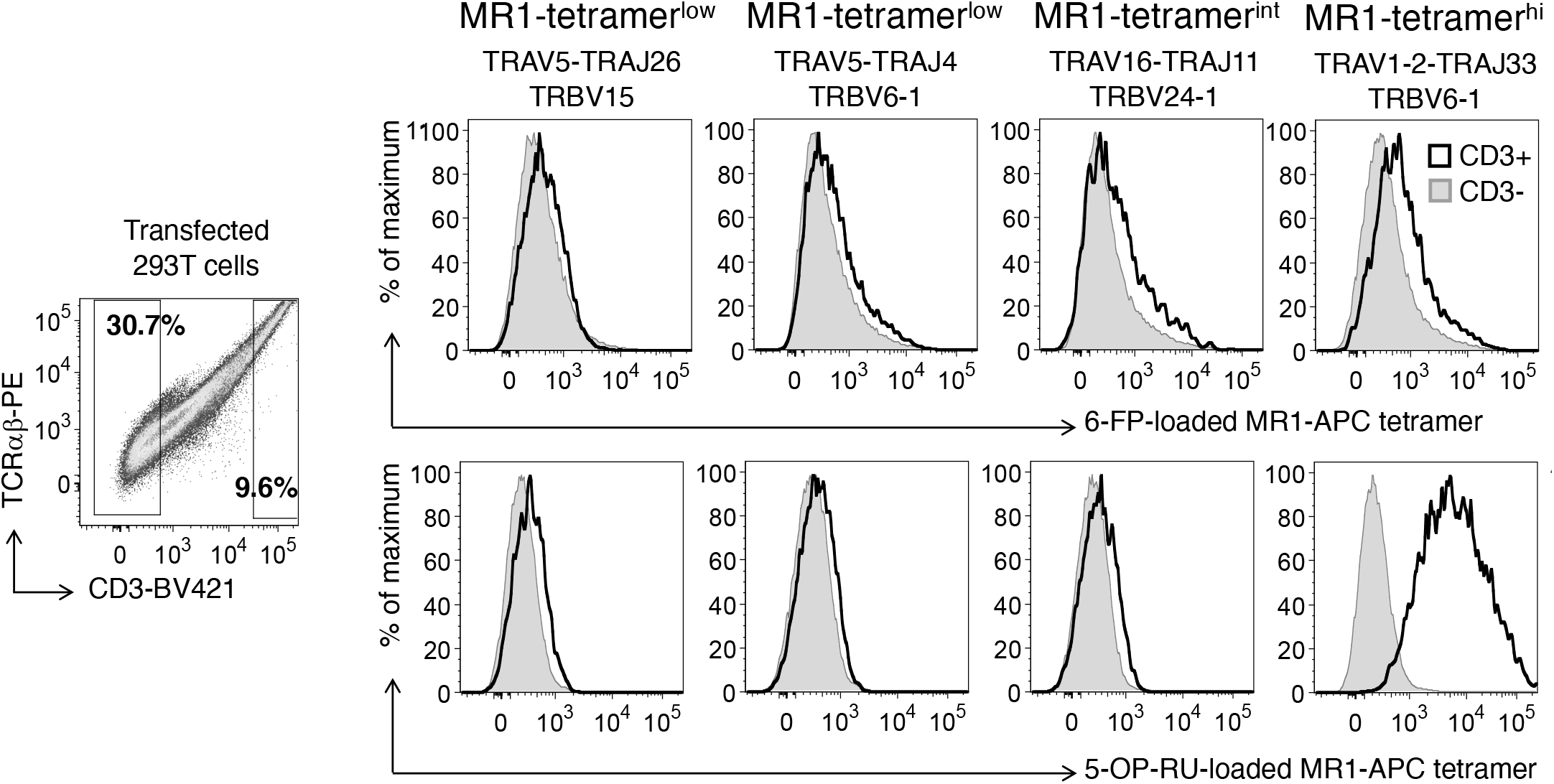
HEK293T cells transfected with non-TRAV1-2 TCRs from MR1-tetramer-sorted cells do not bind MR1. The plots show flow cytometry of human embryonic kidney (HEK) 293-T cells co-transfected with pMIG vectors expressing CD3 and paired TCRα and TCRβ sequences from TCR sequences identified in sorted MR1-tetramer-binding populations with different MR1-binding intensities. The left panel shows gating of CD3- and CD3+ populations used to derive the overlayed histograms are gated on CD3- (grey) and CD3+ (black).

To explain the lack of binding between these TRAV1-2^−^ TCRs and5-OP-RU-loaded MR1, we took a closer look at the TCRβ sequences. Unexpectedly, a single TCRβ sequence consisting of TRBV24-1-TRBJ2-5 with a unique CDR3 nucleotide sequence was detected in 10 out of the 15 TRAV16^+^ single cells **(Supplementary Table 3)**. Interestingly, the same TCRβ nucleotide sequence (TRBV24-1-TRBJ2-5) was paired with the canonical MAIT TCRα TRAV1-2-TRAJ33 in 3 wells **(Supplementary Table 3)**. Because the PCR reactions were performed in multiplex format, we hypothesized that this particular T cell clone expressed two different, functional TCRα chains, but that only one of the PCR products dominated the PCR reaction. Hence, to resolve the discrepancy, we re-amplified the templates that initially gave rise to a TRAV16-TRAJ11 PCR products, using only the TRAV1-specific forward primer, which captures the TCRα variable genes TRAV1-1 and TRAV1-2 only, as previously described^41^. Using this approach, 10 out of the 15 templates initially giving rise to TRAV16-TRAJ11 sequences now gave rise to a PCR product that resulted in identical TRAV1-2-TRAJ33 sequences and paired with the same TRBV24-1-TRBJ2-5 TCRβ **(Supplementary Table 3)**. Whereas we initially interpreted these results as non-canonical TCRs binding to MR1, the data were more consistent with clonal expansion of a T cell co-expressing one TCR β chain, a TRAV1-2^+^ invariant MAIT TCR α chain, and an additional, non-canonical TCRα chain. If only the canonical TCRα chain binds MR1, the lower tetramer binding of these TCRs could be caused by competition of two different TCRα chains with the same TCRβ chain (TRBV24-1-TRBJ2-5), analogous to what has been described for NKT cells^49^.

Finally, to reproduce our finding of dual TCRα expression on MAIT cells in an independent experiment, we analyzed paired TCR sequences in MR1-tetramer-binding cells from different blood donors^55^. Although in this experiment we sorted all MR1 tetramer-binding T cells, including the MR1-tetramer^high^ ones, we identified cells that co-expressed the canonical invariant TRAV1-2^+^ TCR α chain with a TRAV1-2^−^ α chain in PBMC samples from donors of different ages, as well as healthy spleen tissues of deceased donors **(Figure 5)**. Collectively, our study suggests that dual-TCRα expression is common among MR1-tetramer-binding MAIT cells in different human populations, tissue types and disease states.

**Figure 5:**
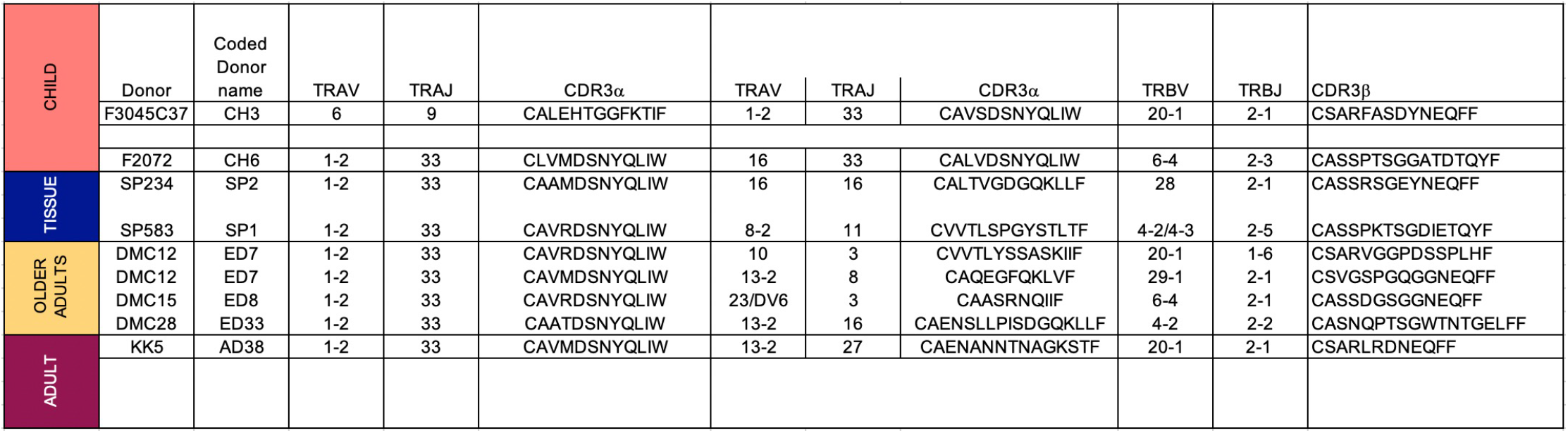
Examples of dual TCRα-expressing MAIT cell clones detected in different sample types. Codes: CH (Child), AD (Adult), ED (Elderly), SP(Spleen).

## Discussion

In this study, we hypothesized that TCRs with decreased affinity for MR1-5-OP-RU would reveal new TCR motifs that may prefer MR1 ligands other than 5-OP-RU or correlate with TB disease. Our hypothesis was motivated by the reported expansion of diverse MAIT cell clonotypes following *Salmonella* challenge of humans in individuals who progress to disease^57^, and the discovery of new antigen classes derived from the related mycobacterium *M. smegmatis*^28^. However, our search for new TCR motifs based on differential binding to the 5-OP-RU-loaded MR1 tetramer was confounded by the co-expression of two TCRα chains in the same T cell. The phenomenon of dual TCRα co-expression has been previously described for MHC-restricted ^58, 59^ and CD1d-restricted^49^ T cell subsets. Unlike the TCRβ locus, the TCRα counterpart is not subject to strict allelic exclusion, so dual TCRα expression is more common^60, 61^. TCRα recombination is also known to occur simultaneously on both alleles to maximize productive TCRαβ recombination and diversity in the TCR repertoire^62^.

The simplest explanation for the lower MR1-tetramer staining, which is also supported by these reports of dual TCRα chains in other systems, is that the canonical MAIT TCR binds to MR1, but the competition of the two TCRα chains to pair with the same pool of available TCRβ chains reduces the MR1-tetramer-binding intensity by reducing functional TCR expression on the cell surface. Hence, the hypothesis that these TCRs displayed preferential affinity to different MR1 antigens was not supported by the data. Importantly, our data point to a potentially common artifact in interpreting TCRα sequences, particularly from high-dimensional sequencing data^63^. Since research focuses on identifying TCR motifs and antigen specificities of non-MHC-restricted DURT cells, including MAIT cells, new TCR motifs require systematic validation for MR1-specificity through TCR transfer, especially in light of the reported low frequency of TRAV1-2^−^ MAIT cells^13, 20, 33, 34^.

We detected dual TCRs or lower tetramer staining in multiple donors studied with different methods in two laboratories. These unexpectedly common observations suggest that T cells with invariant TCRα chains may even have a higher propensity for expression of two TCRα chains compared to conventional MHC-restricted T cells. Several known aspects of conserved TCR gene usage on MAIT cells are consistent with this hypothesis. Firstly, innate T cells, including MAIT^24, 64^, type I NKT cells^65^, and germline-encoded mycolyl lipid-reactive (GEM) T cells^66^, express TCRs that mostly consist of genome-encoded segments, and few N nucleotides^7, 67^. TCRα recombination starts from the proximal Vα and Jα genes, and proceeds outwardly towards distal Vα and Jα segments until a productive rearrangement occurs or the cell undergoes apoptosis^2^. TRAV1-2 is the second most distal TCR Vα gene, located near the 5’ end of the TRA/D locus. The reliance of many invariant T cells on distal TCRα rearrangements involving TRAV1-2 raises the possibility that their thymic progenitors had extended survival windows during the CD4^+^CD8^+^ double positive (DP) thymocyte stage^68^, when TCRα recombination took place. However, this hypothesis warrants additional studies.

Prior reports suggest that dual TCR co-expression, i.e., two TCRα chains sharing a single TCRβ chain, could shelter one TCRαβ pair from thymic negative selection and consequently increase the propensity for autoimmunity, as reported in mice ^48, 69, 70^ and humans^71, 72^. The second reason for a higher likelihood of MAIT and NKT cells to express two TCRα chains is that both subsets have been reported to undergo selection on hematopoietic cells, namely other DP thymocytes, expressing MR1^73, 74, 75^ or CD1d^76^, respectively, as opposed to thymic epithelial cells on which MHC-restricted cells are selected^77^. Therefore, selection of MAIT and NKT cell progenitors in separate niches from where strongly autoreactive MHC-restricted cells are often eliminated may render them more likely to escape negative selection despite the potential autoreactivity of the second TCRαβ receptor.

We restricted the analysis in this study to MR1-tetramer-binding MAIT cells, with the aim of identifying unique MAIT TCR motifs, and potentially novel antigenic specificities, as recently described^20, 28, 33, 34, 35^. To our knowledge, a systematic analysis of the propensities of MHC-restricted T cells and DURTs for expression of dual TCRα chains has not been formally conducted. While our analyses were not intended to directly compare the frequency of dual TCRα expression in donor-unrestricted (innate-like) and MHC-restricted T cells, our study calls for caution when identifying new TCR motifs, particularly in DURTs. These DURTs have unique rules for recognition of non-peptide antigens and antigen-presenting molecules^78^, and hence, functional validation of new TCR motifs is fundamental to this growing field. Collectively, our findings support that TRAV1-2 is the dominant TCRα gene used for recognition of MR1-5-OP-RU, consistent with the reported low frequency of alternative MAIT TCRα V-genes^13, 20, 33^.

## Supporting information

Suliman et al_Dual TCRalpha MAIT cells_Supplementary Tables

## Acknowledgements

This study was supported by the National Institutes of Health (NIH) TB Research Unit Network, Grant U19 AI111224-01. SS received the Justice, Equity, Diversity and Inclusion (JEDI) award, which provided free language editorial service on the manuscript. AJC is supported by a Future Fellowship (FT160100083) from the Australian Research Council, an Investigator Grant from the National Health and Medical Research Council of Australia (1193745) and a Dame Kate Campbell Fellowship from the University of Melbourne. The content is solely the responsibility of the authors and does not necessarily represent the official views of the National Institutes of Health. KK was supported by the NHMRC Leadership Investigator Grant (#1173871).

## Declaration

LKN, AJC, JMcC, and JR are named co-inventors on patents describing MR1 tetramers. The MR1 tetramer technology was developed jointly by Prof. James McCluskey, Prof. Jamie Rossjohn, and Prof. David Fairlie, and the material was produced by the NIH Tetramer Core Facility as permitted to be distributed by the University of Melbourne.

